# NOD1 is super-activated through spatially-selective ubiquitination by the *Salmonella* effector SspH2

**DOI:** 10.1101/2021.10.08.463692

**Authors:** Cole Delyea, Shu Y. Luo, Bradley E. Dubrule, Olivier Julien, Amit P. Bhavsar

## Abstract

As part of its pathogenesis, *Salmonella enterica* serovar Typhimurium delivers effector proteins into host cells. One effector is SspH2, a member of the novel E3 ubiquitin ligase family, interacts with, and enhances, NOD1 pro-inflammatory signaling, though the underlying mechanisms are unclear. Here, we report the novel discovery that SspH2 interacts with multiple members of the NLRC family to enhance pro-inflammatory signaling that results from targeted ubiquitination. We show that SspH2 modulates host innate immunity by interacting with both NOD1 and NOD2 in mammalian epithelial cell culture. We also show that SspH2 specifically interacts with the NBD and LRR domains of NOD1 and super-activates NOD1- and NOD2-mediated cytokine secretion via the NF-κB pathway. Mass spectrometry analyses identified lysine residues in NOD1 that were ubiquitinated after interaction with SspH2. Through NOD1 mutational analyses, we identified four key lysine residues that are required for NOD1 super-activation by SspH2, but not its basal activity. These critical lysine residues are positioned in the same region of NOD1 and define a surface on NOD1 that is targeted by SspH2. Overall, this work provides evidence for post-translational modification of NOD1 by ubiquitin, and uncovers a unique mechanism of spatially-selective ubiquitination to enhance the activation of an archetypal NLR.

**SYNOPSIS:** SspH2 is an E3 ubiquitin ligase injected by *Salmonella* Typhimurium into host cells that induces pro-inflammatory signaling. The immune receptor, NOD1, is ubiquitinated in the presence of SspH2, resulting in increased pro-inflammatory cytokine secretion.

- SspH2 super-activates NOD1 and NOD2 to increase pro-inflammatory cytokine secretion, in part, through the NF-κB pathway
- Ubiquitin modification of NOD1 were identified by mass spectrometry
- A specific region of NOD1 is targeted by SspH2 to enhance NOD1 activity.

## INTRODUCTION

*Salmonella enterica* serovar Typhimurium (*S*. Typhimurium) is a Gram-negative facultative intracellular pathogen. It is a major cause of diarrhoeal disease worldwide, and results in 33 million healthy life years being lost yearly (1). A hallmark of *S*. Typhimurium infection is its ability to induce its uptake into non-phagocytic cells, establish a replicative niche inside the cell and modulate the host immune response (2, 3). *S*. Typhimurium uses two type 3 secretion systems encoded on *Salmonella* pathogenicity islands (SPIs) to inject bacterial effectors into host cells to instigate these hallmark processes (4). Among these effectors are novel E3 ubiquitin ligases (NEL) that interfere with host ubiquitination and subvert host cellular processes (5).

*Salmonella* secreted protein H2 (SspH2) is one of three *S*. Typhimurium NELs that share physical similarities with other bacterial NEL proteins, e.g. IpaH4.5 in *Shigella* spp (6, 7). Functional roles in bacterial pathogenesis have been uncovered for the three NELs in *S*. Typhimurium (8–11). These proteins all share a similar two domain structure – an amino terminal leucine-rich repeat (LRR) domain, and a carboxy terminal E3 ubiquitin ligase domain (6, 7, 12, 13). *Salmonella* virulence is compromised in an animal model in the absence of SspH1 and SspH2 (12). Remarkably, SspH2 causes activation of inflammation in host cells that is dependent on its E3 ligase activity (10). Upon injection via the type 3 secretion system, SspH2 is localized to the host plasma membrane via palmitoylation on Cys9 (14). This enables SspH2 to interact with proximal host proteins at this site.

The recognition of microbial- or damage-associated molecular patterns (MAMPs and DAMPs, respectively) on the surface, or in the cytosolic compartment, of host cells is critical for host immunity. Nod-like receptors (NLRs) are associated with sensing MAMPs alongside DAMPs, and ER stress in mammalian cells (15). NLRs are characterized by their tripartite structure consisting of: i) a carboxy terminal LRR domain that senses bacterial structures, ii) a central nucleotide binding domain (NBD) that facilitates NLR oligomerization, and iii) a variable amino terminal domain, which is critical for interactions with downstream effector proteins (16). NOD1 and NOD2 are the founding members of the NLRs, first described in 1999(17), and have since been grouped as NLRCs as they contain a characteristic amino terminal caspase activation and recruitment domain (CARD). NOD1 and NOD2 are commonly found in the cytosol of host cells (17), but they localize to actin-rich regions in the plasma membrane when activated (18, 19).

NOD1 is ubiquitously expressed in cells and is critical for epithelial cell sensing of primarily intracellular Gram-negative peptidoglycan by binding to γ-D-glutamyl-*meso*-diaminopimelic acid fragments of their cell walls(19, 20). Upon ligand recognition through the LRR domain, NOD1 unfolds and oligomerizes to cause NF-κB activation through homophilic CARD domain interactions with RIP2(17, 21, 22). This leads to the secretion of pro-inflammatory cytokines such as IL-8(15). NOD2, a NOD1 homolog, is found primarily in monocytes(23). NOD2 recognizes bacterial peptidoglycan through muramyl dipeptide(24, 25). After ligand binding to the LRR domain and subsequent unfolding, NOD2 activates in a way similar to NOD1, where the NBD region facilitates oligomerization and the CARD domain interacts with RIP2, leading to activation of NF-κB(23).

Bacterial E3 ligases have been reported to interfere with downstream inflammatory signaling processes, such as IpaH4.5 targeting TBK1 for degradation to prevent inflammation(26). Even though NLRs are critical intracellular sensors of pathogens, the literature on bacterial ubiquitination of NLRs is limited. Our previous studies showed that SspH2 interacts with both NOD1, and its adaptor protein, SGT1(10). This interaction led to ubiquitination of NOD1, causing increased secretion of the chemokine IL-8 (10). Interestingly, *S*. Typhimurium can exploit inflammation in intestinal epithelial cells to compete with the microbiota (27, 28). The molecular details of how SspH2 specifically modifies NOD1, and how this leads to enhanced pro-inflammatory signaling remain to be identified. Furthermore, it remains unclear whether the SspH2 interaction with NOD1 is unique, or if it interacts more broadly with other NLRs.

In this study, we used mammalian epithelial cell culture to gain further insights into the biological and mechanistic interactions between the *S*. Typhimurium effector SspH2 and host NLRCs. We show that SspH2 selectively interacts with the LRR and NBD regions of NOD1. We also report that SspH2 interacts with NOD2, where catalytically active SspH2 drives NOD2 super-activation, analogously to SspH2’s effect on NOD1. Furthermore, SspH2-super-activation of NOD1 and NOD2 pro-inflammatory responses were associated with canonical NF-κB signaling. Moreover, we provide evidence that SspH2 ubiquitinates a specific region of NOD1 to trigger increased pro-inflammatory cytokine secretion, thus identifying a mechanism of spatially-selective ubiquitination to enhance activation of an archetypical NLR.

## METHODS

### Tissue culture

HEK293T (ATCC CRL-3216) and HeLa (ATCC CCL-2) cells were cultured in Dulbecco’s modified eagle medium (DMEM) containing 4500 mg/l glucose, L-glutamine, and sodium bicarbonate (Sigma) supplemented with 10% fetal bovine serum (123483-020; Gibco), 100 U/ml penicillin and 0.1 mg/ml streptomycin (P4333; Sigma) and grown at 37°C and 5% CO_2_.

### Cloning

Domain truncation variants of NOD1 were generated by PCR amplification of NOD1 domains using pcDNA3-NOD1-Flag as a template (both pcDNA3-NOD1-Flag and pCMV2-Flag-NOD2 were kindly provided by Dana Philpott, University of Toronto). Primer sequences can be found in supplementary Table 1. Truncation fragments were cloned into pcDNA3.1 using the *KpnI* and *XhoI* restriction enzymes to replace wild type (WT) NOD1. To create the NOD1 domain mutants the following primer combinations were used. CARD: NOD1_CARD_For1 and NOD1_CARD_Rev2; NBD: NOD1_NBD_For1 and NOD1_NBD_Rev1; LRR: NOD1_LRR_For2 and NOD1_LRR_Rev1; ΔCARD: NOD1_NBD_For1 and NOD1_LRR_Rev1; ΔLRR: NOD1_CARD_For1 and NOD1_NBD_Rev1. ΔNBD was created by separately amplifying the CARD and LRR domains using primer pairs: NOD1_CARD_For1/NOD1_CARD_Rev1 and NOD1_LRR_For1/ NOD1_LRR_Rev1. These two PCR fragments were then combined together by crossover PCR. The lysine variants of NOD1 were generated in the pcDNA3.1 NOD1 background, using the Quikchange II site directed mutagenesis kit (Agilent) according to the outlined protocol from Agilent. All mutagenized plasmids were confirmed by Sanger sequencing. All constructs were propagated in *E. coli* DH5α using standard methods. The SspH2 domain expression constructs were from (10).

### Immunoblotting

Proteins were separated by polyacrylamide gel electrophoresis and transferred to nitrocellulose membranes (BioRad). Membranes were dried, rehydrated with Tris-buffered saline (TBS) and blocked with TBS blocking buffer (927-60001; Li-Cor) before incubation with primary antibody diluted in TBS blocking buffer overnight. Membranes were washed and incubated with secondary antibodies diluted in the same buffer for 1 hour. The antibodies used in this study are: mouse α-FLAG (M2; Sigma) 1:2 500; rabbit α-FLAG (SAB4301135; Sigma) 1:2 500; rat α-HA (clone 3F10; Roche Diagnostics); mouse α-Myc (9E10; Santa Cruz Biotechnology) 1:2 500; goat α-mouse (926-68020; Licor) 1:5 000; goat α-rabbit (925-32211; Licor) 1:5 000; goat α-rat (926-32219; Licor) 1:5 000. Blots were imaged with a Li-Cor Odyssey and Image Studio software.

### Manual immunoprecipitation (IP) assay

HEK293T cells were seeded at 1 × 10^6^ cells per 10 cm dish and transfected when cells reached 60-80% confluency with a total of 6 μg of equivalently proportioned plasmid DNA using JetPrime transfection reagent (Polyplus). Lysates were harvested by washing cells with PBS, and applying lysis buffer (20 mM Tris/HCl, pH 7.5, 150 mM NaCl, and 1% Nonidet P-40 (all from Sigma)) supplemented with EDTA-free protease inhibitor mixture cocktail (Roche). Debris was precipitated (16 000x*g*, 20 minutes) and the supernatant fraction was immunoprecipitated as per the protocol from Invitrogen. In brief, 20μl Dynabeads^™^ Protein G beads (Invitrogen) were coupled to 2.5μg α-HA (high affinity) antibody (Roche) or 5 μg α-FLAG (M2 clone) antibody. Antibody-coupled beads were incubated with samples at 4°C overnight and analyzed by immunoblotting.

### NOD1/ NOD2 functional assays

HeLa cells were seeded at 2.5 × 10^5^ cells/well (6-well dish) or 6 × 10^4^ cells/well (24-well dish) and transfected at 60-80% cell confluency with 1μg or 0.25 μg of DNA, respectively, using JetPrime transfection reagent (Polyplus). Two days post-transfection the media was replaced with DMEM containing 0.5% fetal bovine serum and NOD1 or NOD2 agonist [1 μg/ml C12-iE-DAP (Invivogen)+ 10 ng/ml human interferon gamma (AbD serotec) and 5 μg/ml L-18-MDP (InvivoGen) + 10 ng/ml human interferon gamma, respectively]. Following overnight stimulation, secreted IL-8 was quantified by enzyme-linked immunosorbent assay according to the manufacturer’s specifications (BD-Bioscience). For NF-κB pathway inhibition experiments, 20 μM of Bay 11-7082 (Sigma) was pre-incubated with samples for 45 mins before stimulation.

### Automated immunoprecipitation for Mass spectrometry

Transfected HEK293T cell lysate was prepared as outlined above. Automated IP was performed using the KingFisher Duo Prime Purification System using a modification of the manufacturer’s protocol (publication No. MAN0016198; Thermo Scientific) and executed on BindIt software (v4.0.0.45). Reagents were added to a 96 deep-well plate for parallel processing. In brief, 20 μL of Dynabeads^™^ Protein G beads (Invitrogen; 10003D) were bound to 7 μg of anti-DYKDDDDK antibody (SAB4301135; Sigma) in 200 μl of PBS-T (Phosphate Buffered Saline with 0.02% Tween 20) for 10 min, and subsequently washed with 200 μL PBS-T. The Dynabeads with the bound antibody complex were then incubated with 800 μL of transfected cell lysates (~2.5 mg protein) for 10 min at room temperature, and washed twice with 500 μL PBS. The bound proteins were eluted from Dynabeads using 30 μl of 4x Laemmli buffer (Bio-Rad) without reducing agents and heated to 70°C for 10 min. The eluants were collected from the plate, and diluted with 20 μl ddH2O to run on an SDS-PAGE gel.

### In-gel sample preparation for tandem mass spectrometry (LC-MS/MS)

The eluted IP samples were separated by molecular weight using SDS-PAGE (precast 4-20% Mini-PROTEAN TGX Stain-Free Protein Gels, 10 well, 50 μl; Bio-Rad) at 160 V for 10 min. The proteins in the gel were fixed by incubating with 50% EtOH, 2% phosphoric acid at room temperature for 30 min, then the gel was washed twice with ddH_2_O for 10 min. The gel was subsequently stained by Blue-Silver stain (20% pure ethanol, 10% phosphoric acid, 10% w/v ammonium sulfate, 0.12% w/v Coomassie Blue G-250) at room temperature overnight.

After staining, the gel was washed twice with ddH2O for 20 min. Each lane was cut into four fractions of gel pieces and transferred to a round-bottom 96-well plate to be repetitively destained by 150 μl destaining solution (50 mM ammonium bicarbonate, 50% acetonitrile) in each well at 37°C. The gel pieces were then dried by incubating with acetonitrile at 37°C, rehydrated and reduced with 175 μl of reducing solution (5 mM β-mercaptoethanol, 100 mM ammonium bicarbonate) at 37°C for 30 min, and alkylated with 175 μl of alkylating solution (50 mM iodoacetamide, 100 mM ammonium bicarbonate) at 37°C for 30 min. The gel pieces were washed twice with 175 μl of 100 mM ammonium bicarbonate at 37°C for 10 min, and completely dried by incubating with acetonitrile at 37°C. Proteins in each well were digested with 1 μg of sequencing-grade trypsin (Promega Inc.) in 75 μl of 50 mM ammonium bicarbonate and incubated overnight. Tryptic peptides in the gel pieces were extracted by incubating with 2% acetonitrile, 1% formic acid, then with 50% acetonitrile, 0.5% formic acid, each at 37°C for 1 hour. The extracted peptides were transferred to another round-bottom 96-well plate, dried using a Genevac (EZ-2 plus). Each sample was pre-fractionated into four injections for LC-MS/MS.

### Mass spectrometry analyses

Peptides were separated using a nanoflow-HPLC (Thermo Scientific EASY-nLC 1200 System) coupled to Orbitrap Fusion Lumos Tribrid Mass Spectrometer (Thermo Fisher Scientific). A trap column (5 μm, 100 Å, 100 μm × 2 cm, Acclaim PepMap 100 nanoViper C18; Thermo Fisher Scientific) and an analytical column (2 μm, 100 Å, 50 μm × 15 cm, PepMap RSLC C18; Thermo Fisher Scientific) were used for the reverse phase separation of the peptide mixture. Peptides were eluted over a linear gradient over the course of 90 min from 3.85% to 36.8% acetonitrile in 0.1% formic acid. Data were analyzed using ProteinProspector (v5.22.1) against the concatenated database of the human proteome (SwissProt.2017.11.01), with maximum false discovery rate at 5% for proteins, and 1% for peptides. Search parameters included a maximum of three missed trypsin cleavages, a precursor mass tolerance of 15 ppm, a fragment mass tolerance of 0.8 Da, with the constant modification carbamidomethylation (C), and variable modifications of acetyl (protein N-term), deamidated (N/Q), oxidation (M), and GlyGly (uncleaved K). The maximum number of variable modifications was set to 4.

The LC–MS/MS proteomics data have been deposited in the ProteomeXchange Consortium via the MassIVE partner repository with the dataset identifiers MSV000087693 and MSV000087700. The MS/MS spectra are available using MS-Viewer in ProteinProspector 6.3.1 with the following search keys: anti-FLAG IP performed in HEK293T cell lysates overexpressing NOD1+Ub+EV (bwonnmmgfb), NOD1+Ub+SspH2 C580A (ywbpq3gwof), NOD1+Ub+SspH2 WT (bnfydlbojt) NOD1+EV (8ttoweyamd), NOD1+SspH2 C580A (zvtcjoztu2) and NOD1+SspH2 WT (mggv6qt4m2).

### Statistical analysis

All error bars are representative of SD. NOD1 and NOD2 activation were analyzed in HeLa cells and statistical analyses were determined by non-parametric Student’s *t* test (Mann-Whitney U test). The semi-quantitative mass spectrometry peptide fragment analysis was analyzed by linear regression with 95% confidence intervals (seen as a solid line and dotted line, respectively). NOD1 lysine variant activation in HeLa cells was analyzed by 1-way ANOVA with Dunnett’s multiple comparison test to a control sample (NOD1 + EV or against NOD1 + SspH2). All statistical comparisons were performed using Prism 6.01.

## RESULTS

### NOD1 NBD and LRR domains interact with SspH2

We previously reported that SspH2 interacts with SGT1 and NOD1 (12). However, the mechanistic details of how SspH2 interacts with NOD1 remain unexplored. In HEK 293T cells, we transiently transfected NOD1 single domain constructs [CARD (residues 1-160), NBD (residues 134-584), LRR (residues 697-1053)], or single domain deletion constructs [(ΔCARD (residues 160-1053), ΔNBD (residues 1-160 & 580-1053), and ΔLRR (residues 1-584)] to identify which domains are critical for NOD1 interaction with SspH2 (Fig. 1A). Through reciprocal co-immunoprecipitation, we determined that the NBD and LRR domains of NOD1, but not the CARD domain, interact with SspH2 (Fig. 1B).

**Figure 1.**
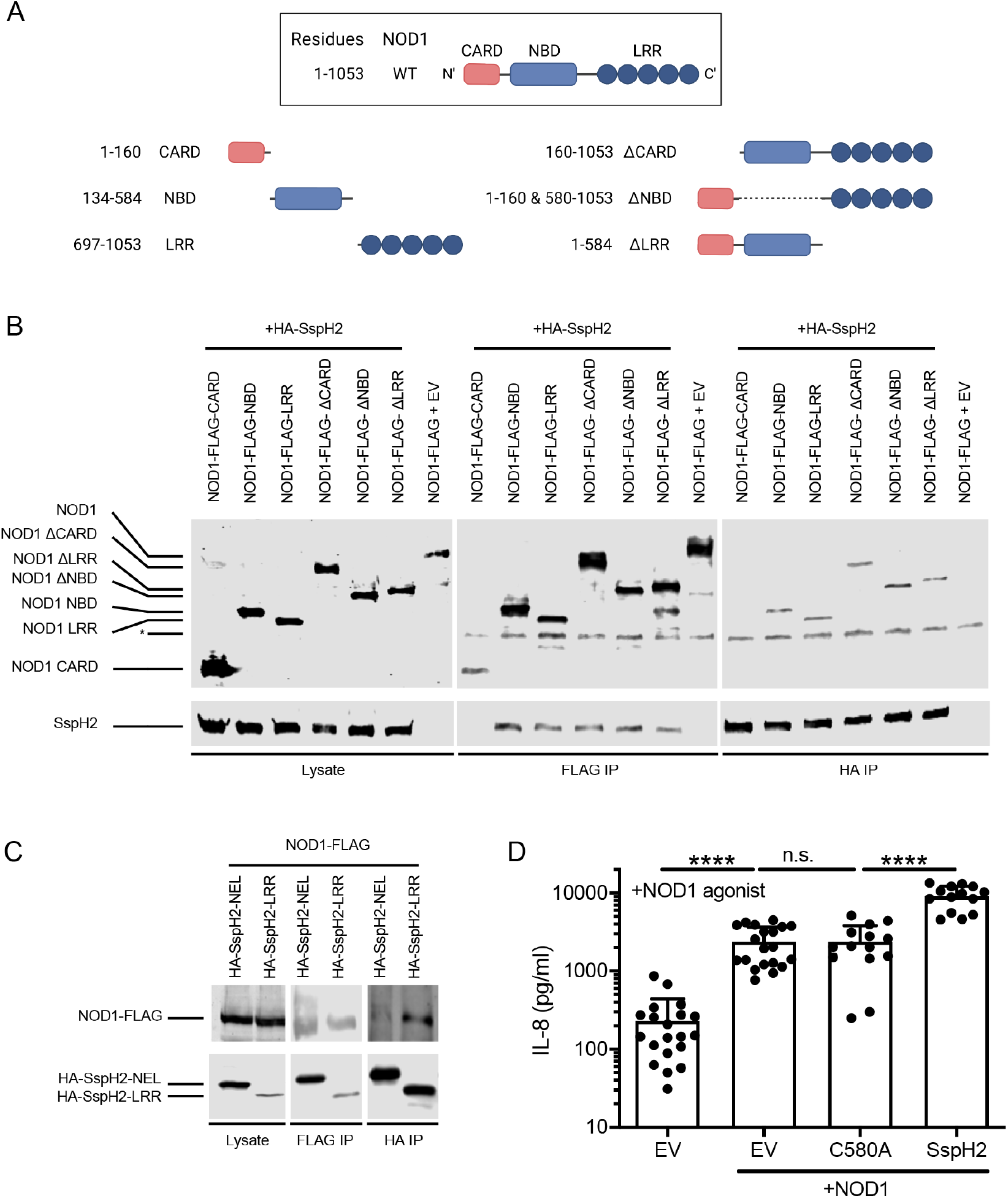
*S*. Typhimurium SspH2 super-activates NOD1 through interactions with the NBD and LRR domains. **A.** Reciprocal co-immunoprecipitation (co-IP) of NOD1 domain fragments with SspH2 transiently expressed in HEK293T cells. * denotes a non-specific protein band. **B.** Reciprocal co-IP analyses of SspH2 domain fragments with NOD1 transiently expressed in HEK293T cells. **C.** IL-8 secretion assay in HeLa cells transiently expressing NOD1, SspH2, SspH2C580A (C580A) or empty vector (EV) as indicated, in the presence of NOD1 agonist (1 μg/mL c12-iE-DAP and 10 ng/mL human IFNγ). IPs and immunoblotting were performed with the indicated antibodies. Data is presented as the mean with standard deviation for 7-9 biological replicates (with 2-3 technical replicates each). Each dot represents 1 technical replicate. Data were analyzed using a non-parametric Mann-Whitney test, **** denote *P* < 0.0001 between the indicated groups.

SspH2 is comprised of a carboxy terminal NEL domain and an amino terminal LRR domain(7). To investigate which domain of SspH2 is responsible for NOD1 binding, we transiently transfected HEK 293T cells with individual SspH2 domains and NOD1. We observed that both the LRR and NEL domains of SspH2 appeared to interact with NOD1 (Fig. 1C). However, we observed that less NOD1 co-immunoprecipitated with SspH2 NEL, compared to SspH2 LRR, indicating that there may be different binding strength between the SspH2 regions (Fig. 1B). We confirmed that SspH2, but not the catalytically inactive C580A variant, induced super-activation of NOD1 by ~4 fold in the presence of NOD1 agonist (Fig. 1D). These data show that NOD1 interacts with SspH2 through its NBD and LRR domains, resulting in super-activation due to the ubiquitin ligase activity of SspH2.

### SspH2 interacts with multiple NLRCs

To ascertain whether SspH2 interaction with NOD1 was unique, we investigated whether it could interact with NOD2, another member of the NLRC family (23). Similar to NOD1, we observed that SspH2 interacts with NOD2 via reciprocal co-immunoprecipitation in cell culture lysate (Fig. 2A). This interaction was specific, as NOD2 did not interact with SspH1 in this assay, despite its 69% sequence homology with SspH2(12). It should be noted that in these experiments, SspH1 levels were lower than SspH2, and thus weak protein interactions between SspH1 and NOD2 could be below the limit of detection. Furthermore, similar to previous observations with NOD1, catalytically inactive SspH2 interacts with NOD2 (Fig. 2B). Again, we observed that both the SspH2 NEL and LRR domains interact with NOD2 (Fig. 2C).

**Figure 2.**
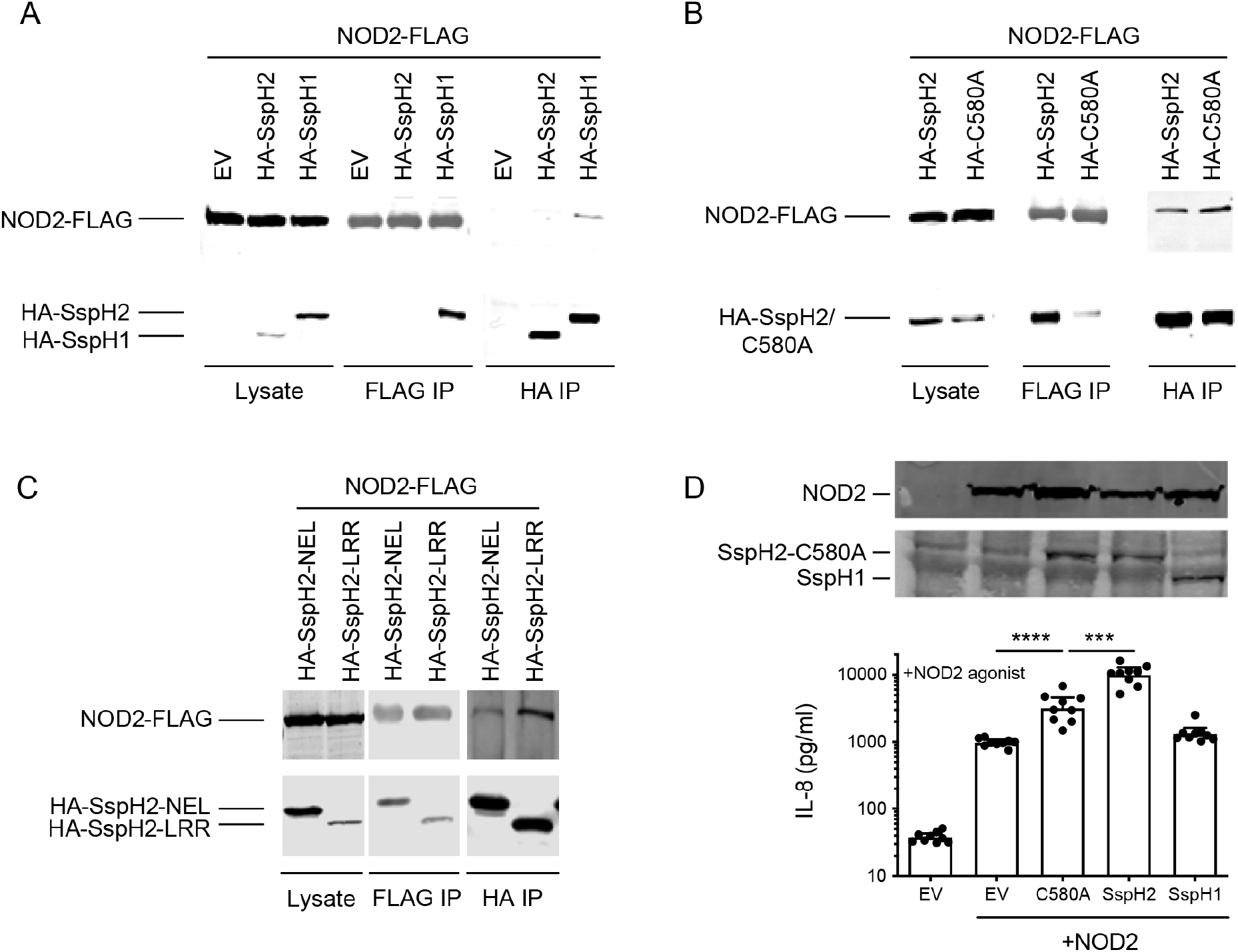
*S*. Typhimurium SspH2 interacts with and super-activates NOD2. **A.** Reciprocal co-IP analysis of NOD2 with SspH2 or SspH1 transiently expressed in HEK293T cells. **B.** Reciprocal co-IP analysis of NOD2 with SspH2 WT or SspH2 C580A transiently expressed in HEK293T cells. **C.** Reciprocal co-IP analyses of SspH2 domain fragments with NOD2 transiently expressed in HEK293T cells. **D.** IL-8 secretion assay in HeLa cells transiently expressing NOD2, SspH2, SspH2C580A (C580A), SspH1, or empty vector (EV) as indicated, in the presence of NOD2 agonist (5μg/mL L-18 MDP and 10ng/mL human IFNγ). Protein expression in HeLa cell lysate following transient expression of indicated constructs. NOD2 was tagged with FLAG. SspH2, SspH2C580A, and SspH1 were tagged with HA. Data is presented as the mean with standard deviation for 3 biological replicates (with 2-3 technical replicates each). Each dot represents 1 technical replicate. Data were analyzed using a non-parametric Mann-Whitney test, *** and **** denote *P* < 0.001 and *P* < 0.0001 respectively between the indicated sample groups.

Having identified a complex between SspH2 and NOD2, we tested if this interaction functionally mimics that of NOD1 and SspH2 in mammalian cell culture in the presence of NOD2 agonist (Fig. 1C). We observed that NOD2 activation was ~10 fold higher in the presence of SspH2. Notably, the presence of SspH2 C580A induced ~3-fold higher activation compared to basal NOD2 activity, whereas NOD2 activity was not increased by SspH1 (Fig 2D). This is consistent with the finding that SspH1 and NOD2 do not interact. We confirmed that all proteins were expressed under our assay conditions (Fig. 2D). We repeated these experiments without NOD2 agonist to determine if NOD2 super-activation by SspH2 was dependent on the NOD2 agonist. Intriguingly, in the absence of NOD2 agonist, SspH2 WT and SspH2 C580A increased IL-8 secretion by ~10 fold and ~7 fold, respectively (Fig. S1A). This result is similar to what has been previously observed with NOD1 (12). Together, this shows that SspH2 interacts with and super-activates NOD2, inducing increased IL-8 secretion, in a similar fashion to NOD1.

In control experiments, we determined the basal effect of SspH2 expression on IL-8 secretion in the presence or absence of NLR agonist without exogenous NLR expression. We observed that SspH2 expression alone increased IL-8 secretion by ~3 fold (Fig. S1B). This was increased to ~10 fold in the presence of NOD1 agonist (Fig. S1C) and NOD2 agonist (Fig. S1D). However, it is worth noting that these levels are still ~20 fold lower than enhancements observed in the presence of exogenous NLR. These data suggest that SspH2 is specifically interacting with host NLRs to cause super-activation and a subsequent increase in IL-8 secretion.

### SspH2-mediated NLR super-activation utilizes NF-κB signaling

NOD1 and NOD2 are thought to produce IL-8 through the activation of the NF-κB pathway (29). To study the downstream effects of SspH2 on NOD1 and NOD2 signaling, IL-8 secretion assays were performed in the presence of the NF-κB pathway inhibitor, Bay 11-7068 (Bay 11). Bay 11 irreversibly inhibits NF-κB activation by blocking the phosphorylation of IκBα, which suppresses the nuclear translocation of p65 and its binding to NF-κB response elements, thus effectively preventing pro-inflammatory cytokine production(30, 31).

As expected, our data showed that the NF-κB inhibitor Bay 11, reduced NOD1 activation as measured by IL-8 secretion. This reduction was ~4.5 fold and ~9 fold in the absence and presence of SspH2 respectively (Fig. 3A). We found that in the presence of Bay11 the levels of IL-8 secretion were comparable, with and without SspH2 (Fig. 3A). We noted a slight decrease in SspH2 levels in the presence of Bay 11; however, the Bay 11 effect is more consistent with NF-κB inhibition because IL-8 secretion levels in Bay 11-treated SspH2 samples were lower than samples that lacked SspH2 (Fig. 3A).

**Figure 3.**
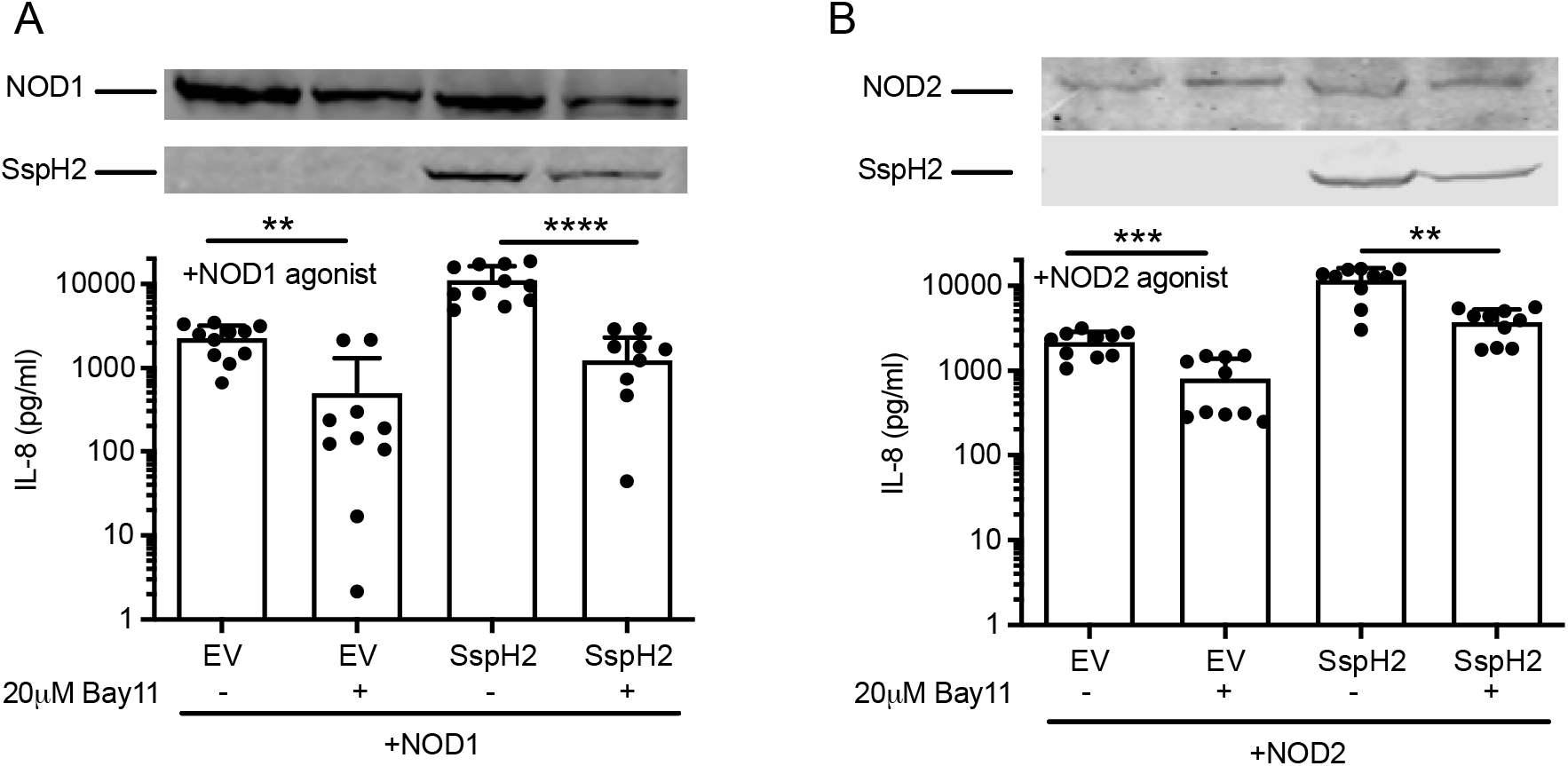
SspH2-mediated NLR super-activation signals through the NF-κB pathway. **A,B.** IL-8 secretion assay in HeLa cells transiently expressing NOD1, SspH2 or EV and treated with NOD1 agonist (1μg/mL C12 iE-DAP and 10ng/mL human IFNγ), with or without the addition of 20μM Bay-11 (**A**); or NOD2, SspH2 or EV and treated with NOD2 agonist (5μg/mL L-18 MDP and 10ng/mL human IFNγ), with or without the addition of NF-κB inhibitor, Bay11 (**B**). NOD1 and NOD2 were tagged with FLAG. SspH2 was tagged with HA. Data is presented as the mean with standard deviation for 4 biological replicates (with 2-3 technical replicates each). Each dot represents 1 technical replicate. Data were analyzed using a non-parametric Mann-Whitney test and **, ***, and **** denote *P* <0.01, < 0.001, < 0.0001 respectively, between the indicated sample groups. See materials and methods for more details.

Due to the similar patterns of protein binding in NOD1 and NOD2, we also investigated the effect of Bay 11 in the NOD2 functional assay. We found that NOD2 signaling was also diminished with or without SspH2 in the presence of Bay-11, albeit to a lesser extent than what was observed with NOD1. This reduction was ~2.5 fold and ~3 fold in the absence and presence of SspH2, respectively (Fig. 3B). We found that upon Bay 11-treatment, levels of IL-8 secreted in the presence of NOD2 + SspH2 was significantly increased by ~4.5 fold compared to NOD2 + EV (Fig. 3B). This indicates that NOD2 super-activation by SspH2 may not be entirely dependent on NF-κB signaling. Again, SspH2 levels were slightly reduced upon Bay 11 treatment, which could partially contribute to the decreased IL-8 secretion observed in this experiment (Fig. 3B). Taken together, these data indicate that SspH2 mediates increased NLR activation through NF-κB signaling downstream of NOD1, and partially downstream of NOD2.

### Unique NOD1 ubiquitination is detected in the presence of SspH2

To further elucidate how SspH2 super-activates NOD1, we used mass spectrometry-based proteomics to identify putative ubiquitination sites on NOD1. As shown in Fig. 4A, we transiently co-expressed FLAG-tagged NOD1, SspH2 (WT, C580A or empty vector) and ubiquitin in HEK293T cells. Cells were harvested, lysed and NOD1 was immunoprecipitated using an a-FLAG antibody. These NOD1-enriched samples were further resolved on SDS-PAGE, in-gel digested with trypsin, and subjected to LC-MS/MS. To identify ubiquitinated lysines, we identified peptides featuring a remnant Gly-Gly motif on lysine side chains (K-ε-GG), which is derived from the carboxy terminus of ubiquitin after trypsin digestion(32).

**Figure 4.**
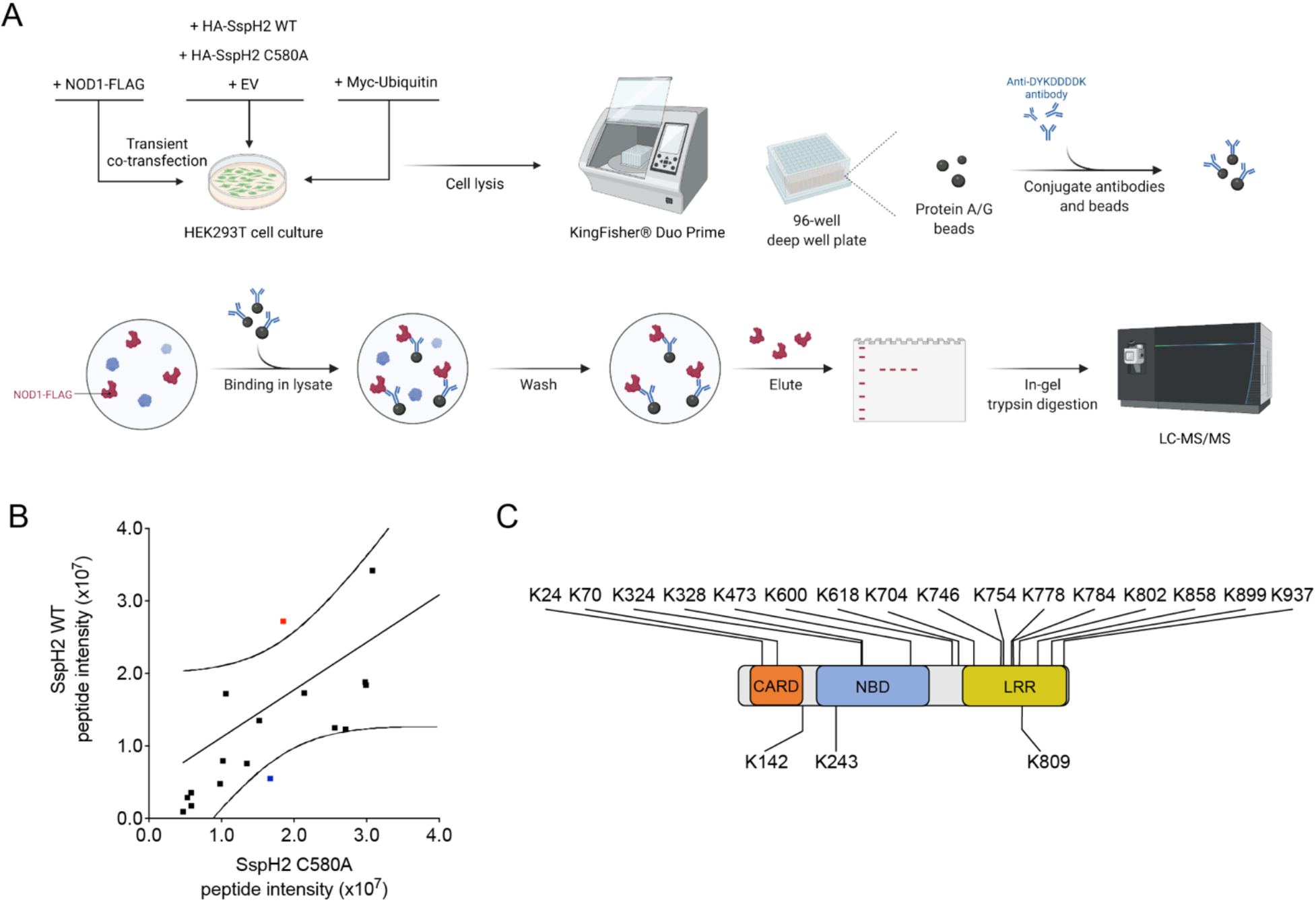
Experimental strategy for identifying NOD1 ubiquitination sites using mass spectrometry. **A.** Schematic diagram of mass spectrometry experiment (created with Biorender). **B.** Gly-gly (Di-glycyl) containing peptide fragment intensity comparison with FLAG-tag immunoprecipitated HEK293T cell lysates overexpressing FLAG-NOD1, HA-SspH2/SspH2 C580A, and Myc-ubiquitin. The solid black line illustrates the linear regression with dotted lines representing the 95% confidence interval. Di-glycyl remnants within the 95% confidence interval are shown in black. Red denotes di-glycyl remnants that were upregulated in SspH2 WT vs SspH2C580A. Blue denotes di-glycyl remnants that were downregulated in SspH2 compared to SspH2C580A. **C.** Schematic representation of all NOD1 lysines changed to arginine in this study.

The overexpressed NOD1 protein was typically detected with more than a hundred peptides with a sequence coverage of ~80%. We found that there were 12 ubiquitination sites in the presence of NOD1 and EV, 23 ubiquitination sites in the presence of SspH2 C580A, and 22 sites on NOD1 in the presence of SspH2. We prioritized candidate lysines for follow-up study if they were unique to SspH2 or showed quantitative differences between samples. For the latter, we performed a semi-quantitative analysis on the intensities of identical peptides present in all of the sample populations. A linear regression was performed on the data utilizing a 95% confidence interval. Data points that fell outside of the linear regression confidence interval were highlighted as candidates of interest. We ascertained first, whether the catalytic activity of SspH2 (WT vs SspH2 C580A) induced significant changes on NOD1 ubiquitination sites (Fig. 4B) and repeated the analysis to compare SspH2 WT or SspH2 C580A against empty vector (Fig. S2A, B). The distribution of lysine residues in NOD1 identified in our mass spectrometry analyses is shown in Fig. S2C. It is noteworthy that to our knowledge, these data provide the first evidence of the location of NOD1 post-translational ubiquitination sites.

Through our analysis, 19 NOD1 lysine residues were selected. 7 were specifically selected for follow-up study because they were unique to SspH2 WT (K142) or fell outside of the linear regression (K328, K473, K776, K778, K784, and K809). The rest were randomly selected to ensure coverage of NOD1 (K24, K70, K324, K600, K618, K704, K746, K754, K802, K858, K899, K937) (Fig. 4C).

### Four lysine residues on NOD1 are critical for its activation by SspH2

To assess whether these prioritized lysine residues were required for basal NOD1 activity, we individually mutated the lysine residues to arginine (to prevent ubiquitination) and tested their activity in our NOD1 functional assay (Fig. 5A). Our data indicated that none of the lysine variants had a suppressive effect on the ability of NOD1 to be activated by its agonist. We did note that the NOD1 variants K784R, K802R and K858R yielded small, but significant increases in basal levels of IL-8 secretion when activated (Fig. 5A). As suggested by the functional assay data, mutation of lysine to arginine did not alter the expression of these NOD1 variant proteins in transiently transfected cells (Fig. S3).

**Figure 5.**
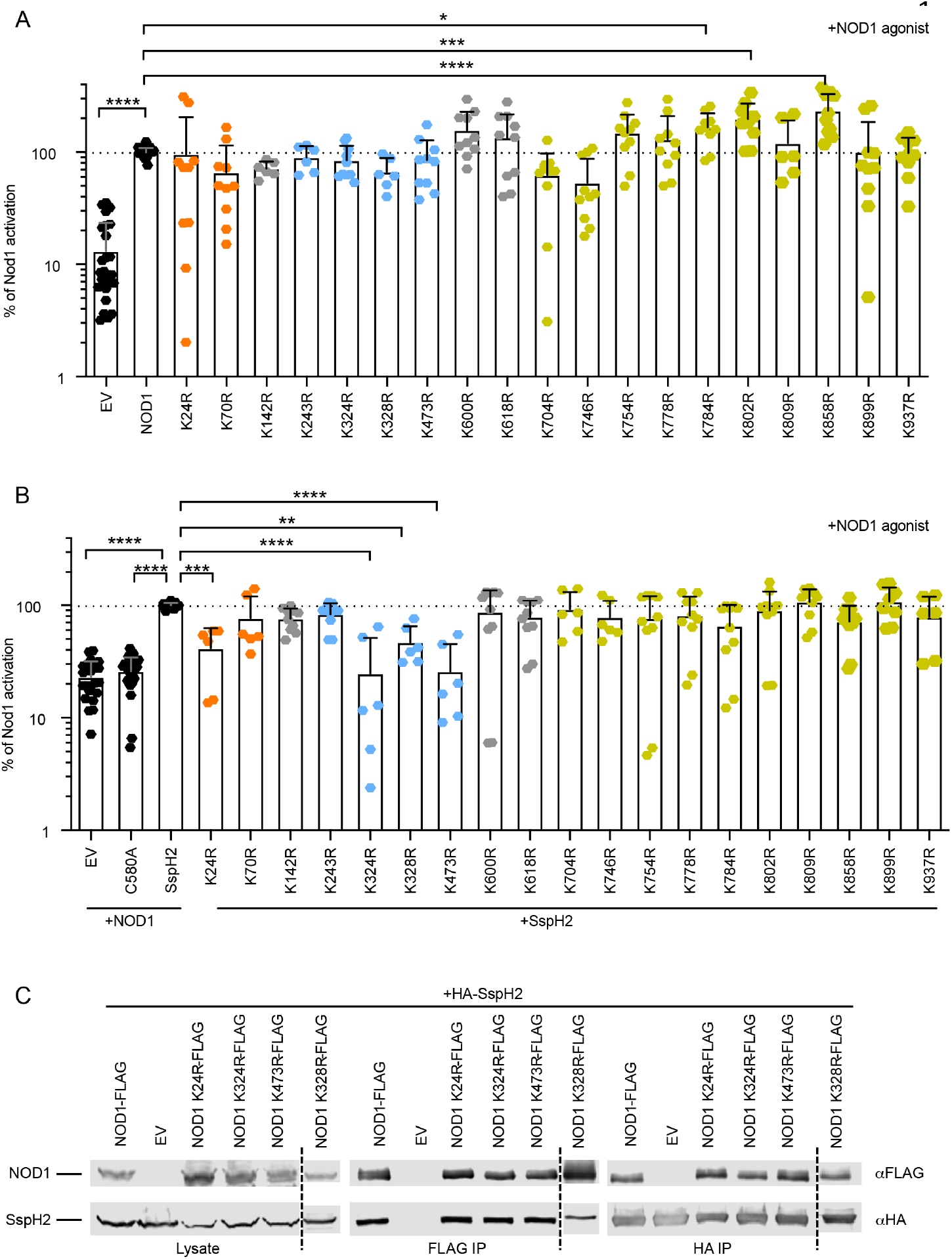
Lysine variants in the NOD1 CARD and NBD domains impact its super-activation by SspH2. **A-B.** IL-8 secretion assay in HeLa cells transiently expressing NOD1, NOD1 lysine variants, C580A, and empty vector (EV) as indicated, in the presence of NOD1 agonist (1μg/mL C12-iE-DAP and 10ng/mL human IFNγ) in the absence (**A**) and presence (**B**) of SspH2. The dotted line represents 100% basal activity (**A**) or super-activation (**B**). **C.** Reciprocal co-IP analysis of NOD1 lysine variants with SspH2 transiently expressed in HEK293T cells. NOD1 lysine variants were tagged with FLAG. IPs and immunoblotting were performed with the indicated antibodies. Data is presented as the mean with standard deviation for 3-5 biological replicates (with 2-3 technical replicates each). Each dot represents 1 technical replicate. Data were analyzed using a One-way ANOVA, *, **, ***, and **** denote *P* < 0.01, *P* < 0.005, *P* < 0.001, and *P* < 0.0001 respectively, between the indicated samples. Dashed line indicates samples run on another gel.

To determine whether any of these lysine residues were important for SspH2 super-activation of NOD1, we repeated the NOD1 functional assay in the presence of SspH2. Intriguingly, lysine to arginine variants in the NOD1 CARD (K24) and NBD (K324, K328, and K473) domains inhibited the SspH2 super-activation phenotype by more than 50% (Fig. 5B). We also noted that K784R, in the LRR region, decreased SspH2 super-activation by ~30% (Fig. 5B).

To ensure that lysine to arginine variants at positions 24, 324, 328 or 473 did not affect the ability of SspH2 to bind to NOD1, we performed reciprocal co-immunoprecipitations from cell lysates co-expressing SspH2 and the variant NOD1 constructs. This analysis indicated that these NOD1 variant proteins still interact with SspH2 (Fig. 5C). Taken together, these data further support a model where SspH2 specifically ubiquitinates lysines in the CARD and NBD domains of NOD1 to augment its pro-inflammatory signaling.

### Lysine residues on one surface of NOD1 are targeted by SspH2 for super-activation

A theoretical protein structure of NOD1 predicted to be of high accuracy by artificial intelligence was recently made available (33). In this model, the lysines that were identified through mass spectrometry are coloured (Fig. 6). Three lysines highlighted in our study, K324, K328, and K473 are present in the same region, where K324 and K473 are both solvent exposed, and K328 is buried (Fig. 6; Fig. S4). This spatial orientation of lysines that are required for NOD1 super-activation is particularly intriguing and supports a structure-function relationship for SspH2 activity.

**Figure 6.**
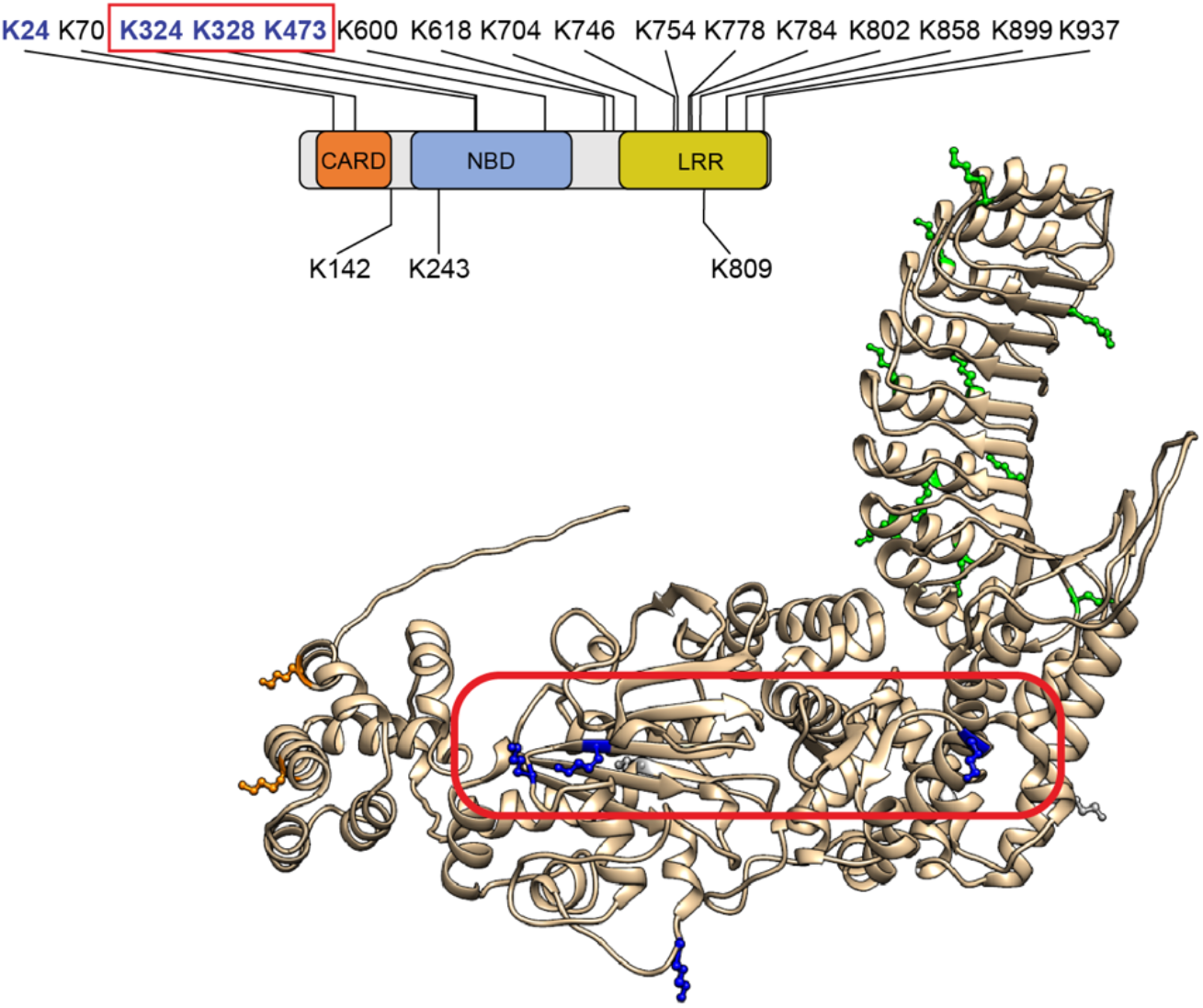
Lysines required for NOD1 super-activation are spatially localized to the same region of NOD1. Conceptual domain model and theoretical structure of NOD1 using the AlphaFold modeling system. The coloured amino acids are lysines found throughout NOD1. Lysine colour correlates to domain location: yellow (CARD), blue (NBD), and green (LRR). Outlined in red are the position of lysines, whose mutation reduces NOD1 super-activation by SspH2 that are on the same surface.

## DISCUSSION

In this study, we uncovered novel aspects of bacterial effector biology through its interaction with an archetypal NLRC. Here, we report that the interaction between the *S*. Typhimurium E3 ubiquitin ligase SspH2 and NOD1 is facilitated by the NBD and LRR domains of NOD1. This interaction leads to SspH2-mediated super-activation of NOD1 via ubiquitination of a specific NOD1 surface.

SspH2 also interacts with, and super-activates, NOD2, although there was residual super-activation by the SspH2 catalytic mutant. The super-activation of NLR activity by SspH2 signals through the NF-κB pathway, although other signaling pathways may make important contributions downstream of NOD2. This difference could possibly be due to differences in protein structure – NOD2 has an additional CARD motif (23), but there may also be different protein interaction patterns or ubiquitination of NOD2. Our ubiquitination analyses focused on NOD1; thus, we do not yet know where SspH2 ubiquitinates NOD2.

Due to the ability of SspH2 to interact with multiple NLRCs, it is tempting to speculate that it interacts with other proteins within this family, or even across NLR families e.g., NLRPs. NLRP3 inflammasome activity is regulated by ubiquitination. Ubiquitination via Pellino2 (34), and de-ubiquitination on the NLRP3 LRR domain via BRCC3 (35, 36) have been shown to activate NLRP3. Alternatively, de-ubiquitination via IRAK1 (34), and ubiquitination by TRIM31 have also been shown to decrease NLRP3 activity by inducing proteasomal degradation in the case of the latter (37). Notably, the bacterial E3 ligase, YopM, has been observed to decrease NLRP3 activation via K63-linked ubiquitination of NLRP3 (38). YopM is an unconventional E3 ubiquitin ligase, in that it contains an LRR domain, but it does not contain a NEL domain (39). Here, we make the novel discovery that SspH2, a *bona fide* member of the NEL family, enhances NLRC pro-inflammatory signaling via targeted ubiquitination.

In our experimental workflow, we anticipated detecting enhanced or differential ubiquitination of NOD1 in the presence of catalytically active SspH2. That we did not observe a significant increase in the number of unique NOD1 ubiquitination sites in the presence of SspH2 suggests that its activity is nuanced. Due to the nature of trypsin cleavage of ubiquitin from lysine residues, we were unable to discern whether this ubiquitination was poly- or mono-ubiquitinated. It has been reported that SspH2 creates K48-linked polyubiquitin chains, suggesting that its targets are meant for proteosomal degradation(40). However, when identifying ubiquitin via immunoblot analysis from samples with ubiquitin and NLR, we did not observe the characteristic smear of poly-ubiquitination (Fig. S5), nor a decrease in protein signal, consistent with our previously reported findings (10). Our analysis of LC-MS/MS peptide intensities revealed that the majority of the ubiquitinated lysines with differential intensities between SspH2 WT and C580A occurred in the LRR domain of NOD1. Surprisingly, we found that mutation of LRR lysines had little effect on SspH2’s capacity to increase pro-inflammatory production, in contrast to lysines in the CARD and NBD regions.

The spatial localization of lysines critical for NOD1 super-activation could suggest that this region of the NBD is where the catalytic cysteine of SspH2 is oriented, or more generally, where the NEL region of SspH2 binds to NOD1. Structural studies could shed further light on these possibilities and contribute to a greater understanding of how NELs interact with NLRs.

There are reports that NOD1 can interact with ubiquitin at Y88 and E84, and at the corresponding sites on NOD2, I104 and L200, in the CARD regions of both proteins (41, 42). This binding of ubiquitin is distinct from post-translational ubiquitin modification, and prevents RIP2 binding to NLRs, to negatively regulate signaling (41, 42). There are also predictions of ubiquitin binding at K436 and K445 on NOD2 (41). It is notable that while there are reports that NLR signaling can be regulated by ubiquitination, there have been no reports of direct ubiquitination on NLRCs. NOD1 and NOD2 interact with host proteins to mediate downstream activation of NF-κB through homotypic CARD domain interaction with RIP2 (17, 22, 23, 43). This process involves many proteins interacting with RIP2 to increase or decrease signaling capacity to maintain appropriate levels of inflammatory activation. Polyubiquitination of RIP2 by host E3 ligases, *e.g*. XIAP, cIAP1/2, ITCH, and Pellino3 induces RIP2 activation upon NOD1/NOD2 stimulation (22, 23, 43). The deubiquitinase proteins A20, OTULIN, and CYLD remove ubiquitin from RIP2 to repress its activation (35, 41, 44). Once RIP2 is ubiquitinated, TAK1 is recruited to phosphorylate IkBa of the IKK complex, leading to NF-κB translocation to the nucleus and production of pro-inflammatory cytokines (45). These conserved host signaling pathways downstream of receptors are where bacterial E3 ligases like IpaH4.5 and SspH1 usually modulate inflammation (11, 26).

To our knowledge, this is the first report that mechanistically links NEL effector activity to functional enhancement of an NLR. Though our cell culture studies do not unequivocally demonstrate that SspH2 directly ubiquitinates NOD1, the report that YopM directly ubiquitinates NLRP3 (38) suggests that SspH2 might also work in a direct fashion. When NOD1 is activated, it undergoes a structural change where the LRR domain is repositioned to relieve its blockade of the CARD and NBD regions (22). It remains to be seen what ubiquitination of NOD1 does to physically alter NOD1 and its ability to interact with other proteins in the cell. One possibility is that ubiquitination of NOD1 leads to structural perturbations that relieve repression by the LRR domain, allowing more facile NOD1 oligomerization. Another possibility is that ubiquitinating NOD1 in the vicinity of adapter binding sites could alter interaction dynamics with RIP2, leading to increased IL-8 secretion (Fig. 7). Our previous observation that SspH2 enhanced NOD1 activity in the absence of NOD1 agonist would be consistent with the first scenario (10), but further study is required to discern between these models. Nevertheless, our current work has illustrated the sophisticated nature of bacterial pathogenesis and uncovered a mechanism whereby a traditionally antimicrobial pathway is subverted for pathogenesis.

**Figure 7.**
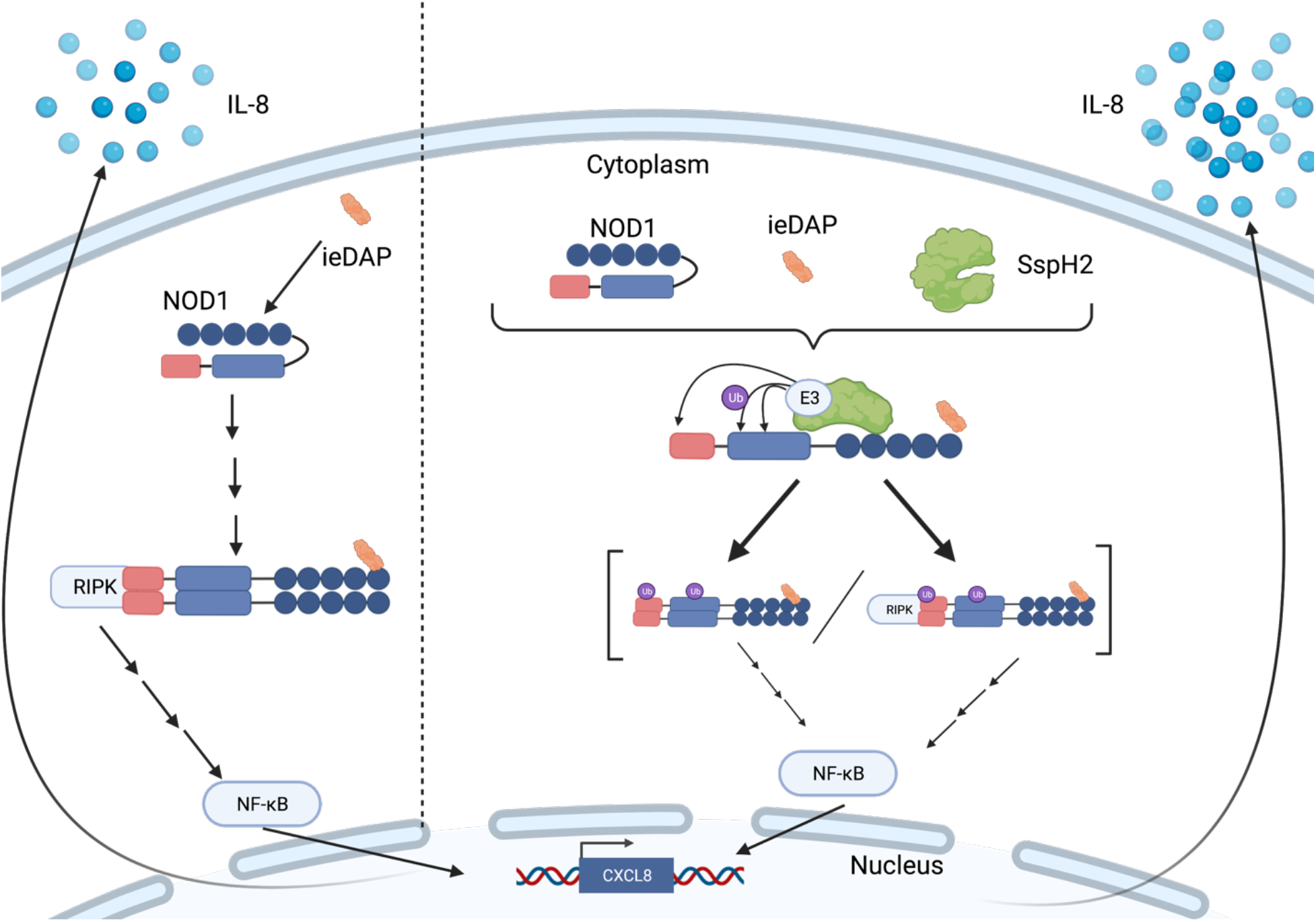
Model of NOD1 super-activation by SspH2 catalyzed ubiquitination. **Left**. In the absence of SspH2, NOD1 activation initiates by interaction with C-12-iE-DAP, unfolding, oligomerizing, and recruiting RIP2 to initiate NF-κB signaling and IL-8 secretion. **Right.** NOD1 super-activation in the presence of SspH2. SspH2 interacts with the NBD and LRR domains of NOD1, and ubiquitinates lysines on the same face of the NBD domain to super-activate NOD1 and cause increased IL-8 secretion. SspH2 ubiquitination of NOD1 may drive its super-activation in several ways including: enhanced oligomerization of NOD1 (left) and augmented interaction dynamics with RIP2 (right).

## ACKNOWLEDGEMENTS

This work was supported by operating grants from the Natural Sciences and Engineering Research Council of Canada (RGPIN-2020-04359 to APB; RGPIN-2018-05881 to OJ) and the Government of Alberta Major Innovation Fund for OneHealth: Antimicrobial Research Consortium (to APB). Infrastructure support was provided by the Canada Foundation for Innovation (37833 and 39051 to OJ). This research has also been funded by the Li Ka Shing Institute of Virology (LKSIoV). CD was supported by studentships from the University of Alberta Faculty of Medicine & Dentistry and LKSIoV. APB holds a Canada Research Chair (Tier 2) in Functional Genomic Medicine and this research was undertaken, in part, thanks to funding from the Canada Research Chairs Program (231622).

## Notes

https://massive.ucsd.edu/ProteoSAFe/static/massive.jsp

### Competing Interest Statement

The authors have declared no competing interest.

### Summary of Updates

Author information updated. Figures 1, 2, 4 and 6 revised.

