## Supplemental Figures 1-5; Supplemental Table 1 for "NOD1 is super-activated through spatially-selective ubiquitination by the *Salmonella* effector SspH2"

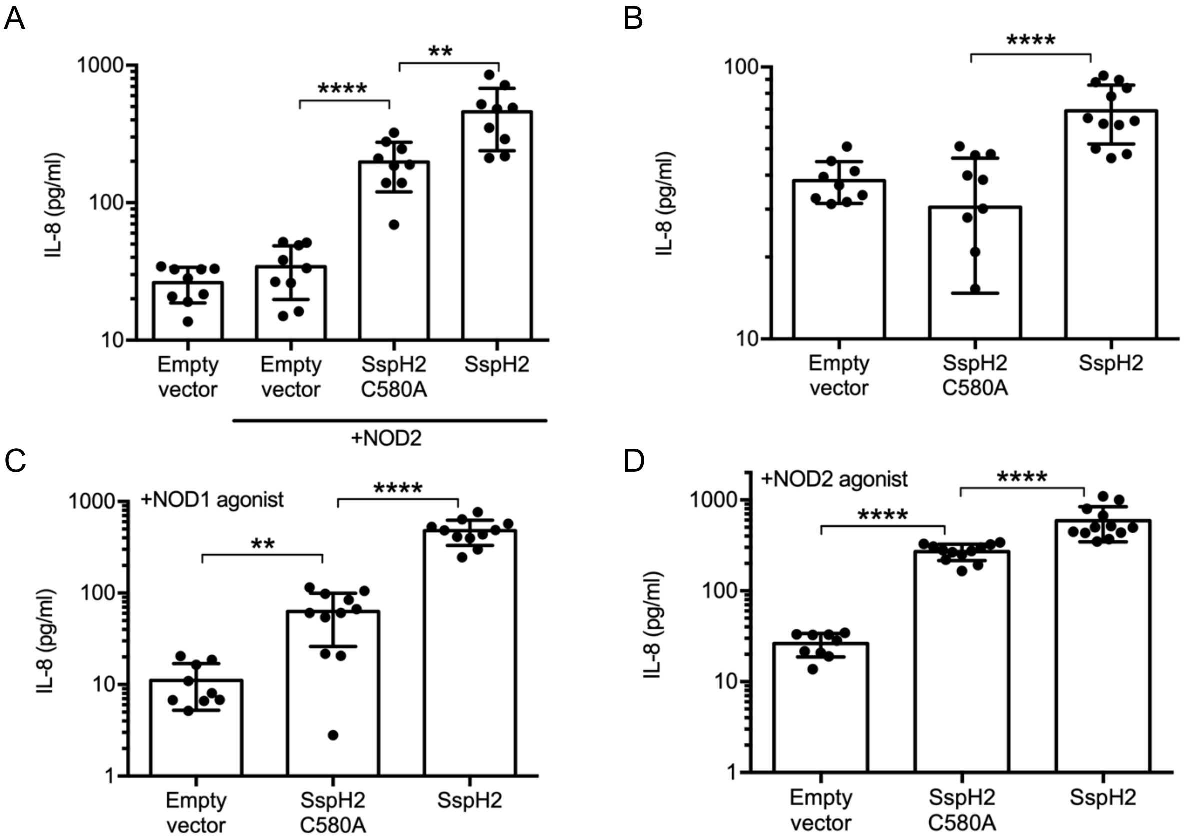


**Supplementary Figure 1. SspH2 can induce IL-8 secretion in the absence of exogenous NLR or its agonist. A**. IL-8 secretion assay in HeLa cells transiently expressing NOD2, SspH2, SspH2C580A (C580A), or empty vector (EV) as indicated, in the absence of NOD2 agonist. **B-D.** SspH2, C580A, and EV as indicated, but not NLRs, in the absence of agonist (**B**), presence of NOD1 agonist (1µg/mL cDAP and 10ng/mL human IFNγ) (**C**), and presence of NOD2 agonist (5µg/mL MDP and 10ng/mL human IFNγ) (**D**). Data are presented as the mean with standard deviation for 3-4 biological replicates (with 2-3 technical replicates each). Each dot represents 1 technical replicate. Data were analyzed using a non-parametric Mann-Whitney test and ** and **** denote $P < 0.01$, $P < 0.0001$ respectively between the indicated sample groups. See materials and methods for more detail.

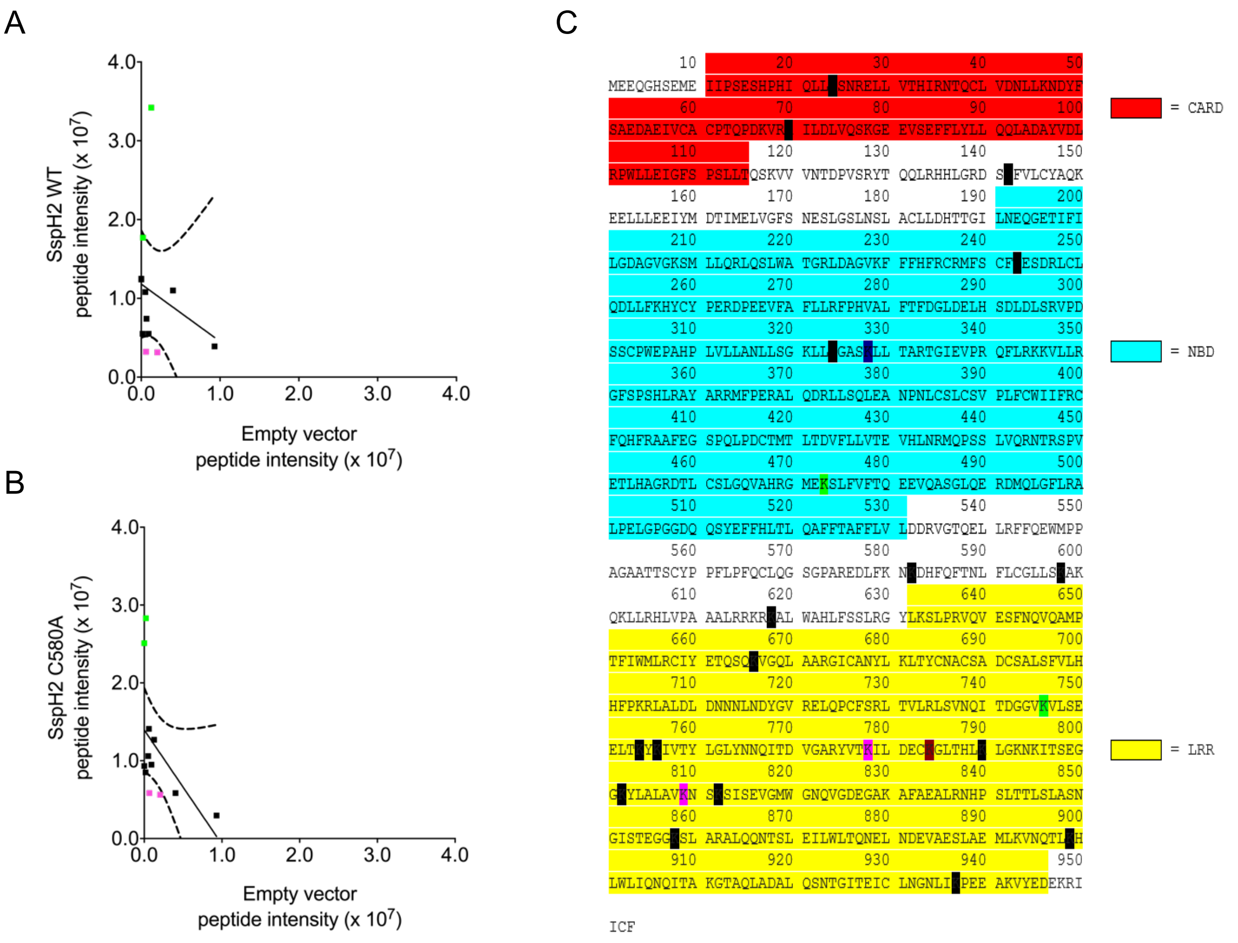


**Supplementary Figure 2. Semi-quantitative analysis of diglycyl peptide intensity identifies NOD1 sites differentially ubiquitinated by SspH2.** Gly-gly (Di-glycyl) containing peptide fragment intensity comparison with immunoprecipitations performed with lysates of HEK 293T cells transiently expressing NOD1, empty vector (EV)/ SspH2 C580A/ SspH2 with ubiquitin. Intensities are scaled on the plot to $1x10^7$. **A.** Di-glycyl modified peptide fragment intensity comparison between SspH2 and EV. **B.** Di-glycyl modified peptide fragment intensity comparison between SspH2 C580A di-glycyl and EV. In each plot the solid black line represents the linear regression with the dotted line indicating the 95% confidence interval. Light green denotes di-glycyl residues that were upregulated in both SspH2/ SspH2 C580A over empty vector (EV). Pink denotes di-glycyl residues that were downregulated in both SspH2/ SspH2 C580A over EV. **C.** Amino acid sequence of NOD1 with Caspase Activation Recruitment Domain (CARD), Nucleotide Binding Domain (NBD), and Leucine Rich Repeat (LRR) regions highlighted. Individual lysine residues highlighted in black identify lysine residues that were identified with di-glycyl modifications. Red denotes di-glycyl residues that were upregulated in SspH2 vs SspH2 C580A. Blue denotes residues that were downregulated in SspH2 compared to SspH2 C580A. Light green denotes di-glycyl residues that were upregulated in both SspH2/ SspH2 C580A over empty vector (EV). Pink denotes di-glycyl residues downregulated in both SspH2/ SspH2 C580A over EV.

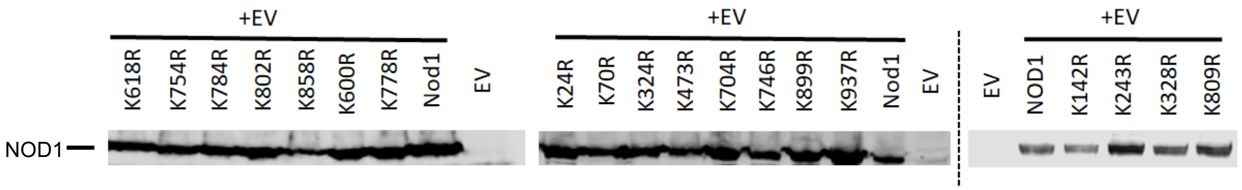


**Supplementary Figure 3. NOD1 lysine variants are stably expressed in vitro.** Protein levels of NOD1 lysine variants or empty vector (EV) transiently expressed in HEK293T cells. NOD1 lysine variants were tagged with FLAG. Immunoblotting was performed with FLAG antibodies. Dashed line indicates samples run on another gel.

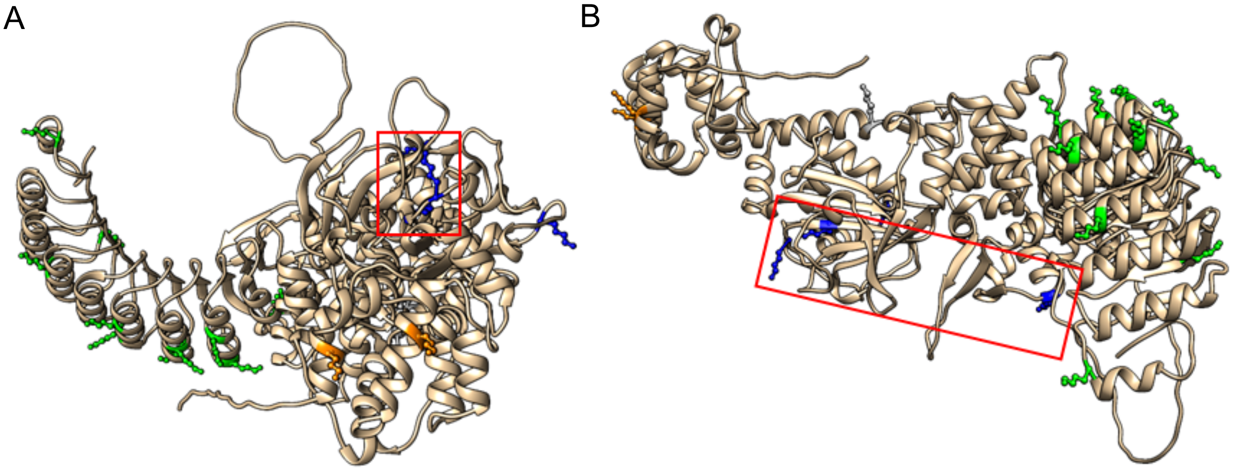


**Supplementary Figure 4. Multiple angles of spatial positioning of lysine variants across NOD1.** Conceptual domain model and theoretical structure of NOD1 using the AlphaFold modeling system (33). NOD1 is viewed from **A.** the left (with the CARD region in front) and **B.** the top-down. The coloured amino acids are lysines found throughout NOD1. Lysine colour correlates to domain location: yellow (CARD), blue (NBD), and green (LRR). Outlined in red are the position of lysines, whose mutation reduces NOD1 super-activation by SspH2 that are in the same surface.

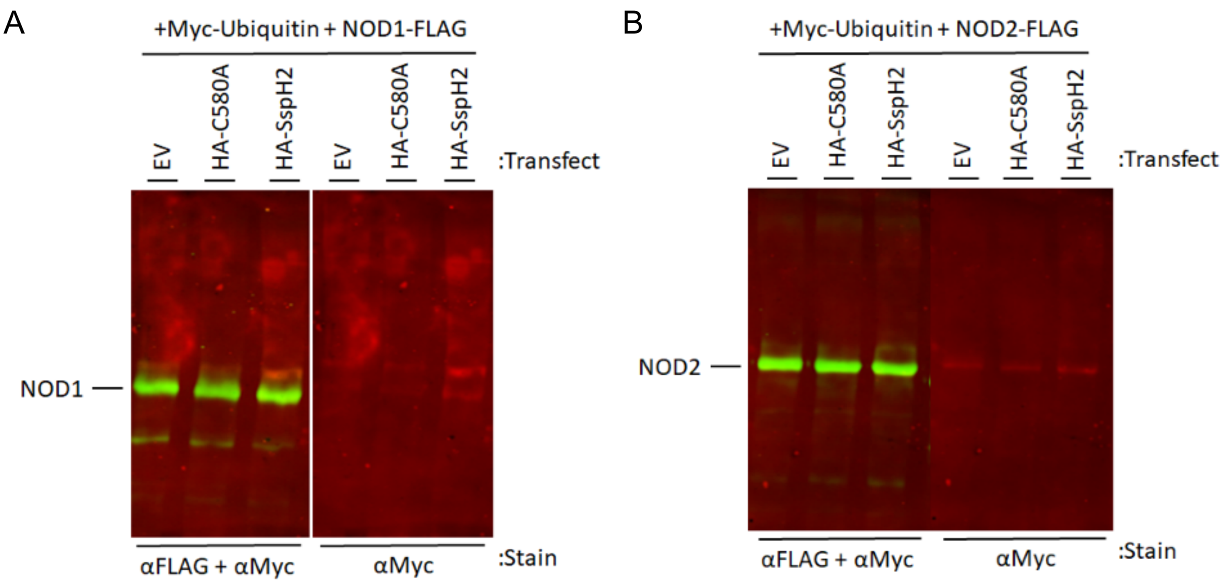


**Supplementary Figure 5. SspH2 does not polyubiquitinate NLRs.** Immunoprecipitation analysis of NLRs transiently transfected in HEK 293T cells. NLRs were tagged with FLAG antibody, and FLAG IP was performed. All samples had Myc-tagged ubiquitin and either EV, SspH2C580A or SspH2 transiently transfected into them with either **A.** NOD1 or **B.** NOD2. Immunoblotting was performed with indicated antibodies. Data is representative of 3 biological replicates.

Supplementary Table 1. Primers used for cloning

| Primer name | Nucleotide sequence |
|---|---|
| NOD1_a71g | 5'-ccccacattcaattactgagaagcaatcgggaacttctg-3' |
| NOD1_a71g_antisense | 5'-cagaagttcccgattgcttctcagtaattgaatgtgggg-3' |
| NOD1_a209g | 5'-cctgacaaggtccgcagaattctggacctggta-3' |
| NOD1_a209g_antisense | 5'-taccaggtccagaattctgcggaccttgtcagg-3' |
| NOD1_a425g | 5'-ggccgtgactccaggttcgtgctgtgc-3' |
| NOD1_a425g_antisense | 5'-gcacagcacgaacctggagtcacggcc-3' |
| NOD1_a728g | 5'-gcatgttcagctgcttcagggaaagtgacaggc-3' |
| NOD1_a728g_antisense | 5'-gcctgtcactttccctgaagcagctgaacatgc-3' |
| NOD1_a971g | 5'-cagtgggaagctgctcagggggctag-3' |
| NOD1_a971g_antisense | 5'-ctagccccctgagcagcttcccactg-3' |
| NOD1_a983g | 5'-ctcaagggggctagcaggctgctcacagc -3' |
| NOD1_a983g_antisense | 5'-gctgtgagcagcctgctagcccccttgag -3' |
| NOD1_a1418g | 5'-ccggggcatggagaggagcctctttgtct-3' |
| NOD1_a1418g_antisense | 5'-agacaaagaggctcctctccatgccccgg-3' |
| NOD1_a1745g | 5'-cgggaagacctcttcaagaacagggatcacttcca-3' |
| NOD1_a1745g_antisense | 5'-tggaagtgatccctgttcttgaagaggtcttcccg-3' |
| NOD1_a1799g | 5'-ggctgttgtccaaagccagacagaaactcctgcg-3' |
| NOD1_a1799g_antisense | 5'-cgcaggagtttctgtctggctttggacaacagcc-3' |
| NOD1_a1853g | 5'-cctgaggagaaagcgcagggccctgtg-3' |
| NOD1_a1853g_antisense | 5'-cacagggccctgcgctttctcctcagg-3' |
| NOD1_a2111g | 5'-cctgcatcacttccccaggcggctggc-3' |
| NOD1_a2111g_antisense | 5'-gccagccgcctggggaagtgatgcagg-3' |
| NOD1_a2237g | 5'-cactgacggtggggtaagggtgctaagcg-3' |
| NOD1_a2237g_antisense | 5'-cgcttagcacccttaccccaccgtcagtg-3' |
| NOD1_a2261g | 5'-gctaagcgaagagctgaccagatacaaaattgtgacctatt-3' |
| NOD1_a2261g_antisense | 5'-aataggtcacaattttgtatctggtcagctcttcgcttagc-3' |
| NOD1_a2333g | 5'-gccaggtacgtcaccagaatcctggatgaatgc-3' |
| NOD1_a2333g_antisense | 5'-gcattcatccaggattctggtgacgtacctggc-3' |
| NOD1_a2351g | 5'-ccaaaatcctggatgaatgcagaggcctcacgc-3' |
| NOD1_a2351g_antisense | 5'-gcgtgaggcctctgcattcatccaggattttgg-3' |
| NOD1_a2405g | 5'-caagtgaaggagggaggtatctcgccctggc-3' |
| NOD1_a2405g_antisense | 5'-gccagggcgagatacctccctccttcacttg-3' |
| NOD1_a2426g | 5'-ctcgccctggctgtgaggaacagcaaatcaatct-3' |
| NOD1_a2426g_antisense | 5'-agattgatttgctgttcctcacagccagggcgag-3' |
| NOD1_a2696g | 5'-gaaatgttgaaagtcaaccagacgttaaggcatttatggcttatc-3' |

| | |
|---|---|
| NOD1_a2696g_antisense | 5'-gataagccataaatgccttaacgtctggttgactttcaacatttc-3' |
| NOD1_a2810g | 5'-ttgcctaaatggaaacctgataagaccagaggaggc-3' |
| NOD1_a2810g_antisense | 5'-gcctcctctggtcttatcaggtttccatttaggcaa-3' |
| NOD1_CARD_For1 | 5’-gaggggtaccgatatcaccatgg-3’ |
| NOD1_CARD_Rev1 | 5’-ctggaagtgatccttgttcatgtagatctcctccag-3’ |
| NOD1_CARD_Rev2 | 5’-gaggctcgagcatgtagatctcctccag-3’ |
| NOD1_NBD_For1 | 5’-gaggggtaccgatatcaccatggacaccatcatggagctggttggc-3’ |
| NOD1_NBD_Rev1 | 5’-gaggctcgagcttgaagaggtcttcccg-3’ |
| NOD1_LRR_For1 | 5’-ctggaggacgatctacatgaacaacaaggatcacttccag-3’ |
| NOD1_LRR_For2 | 5’-gaggggtaccgatatcaccatgaacaaggatcacttccag-3’ |
| NOD1_LRR_Rev1 | 5’-gaggctcgaggaaacagataatccg-3’ |
